# Split minireporters facilitate monitoring of gene expression and peptide production in linear cell-free transcription-translation systems

**DOI:** 10.1101/2024.05.16.594532

**Authors:** Antoine Lévrier, Julien Capin, Pauline Mayonove, Ioannis-Ilie Karpathakis, Peter Voyvodic, Angelique DeVisch, Ana Zuniga, Martin Cohen-Gonsaud, Stéphanie Cabantous, Vincent Noireaux, Jerome Bonnet

## Abstract

Cell-free transcription-translation (TXTL) systems expressing genes from linear dsDNA enable rapid prototyping of genetic devices while avoiding cloning steps. However, repetitive inclusion of a reporter gene is an incompressible cost and sometimes accounts for most of the synthesized DNA length. Here we present minireporters based on split-GFP systems that reassemble into functional fluorescent proteins and can be used to monitor gene expression in *E. coli* TXTL. The 135 bp GFP10-11 fragment produces a fluorescent signal comparable to its full-length GFP counterpart when reassembling with its complementary protein synthesized from the 535 bp fragment expressed in TXTL. We show that minireporters can be used to characterize promoter libraries, with data qualitatively comparable to full-length GFP, and matching with *in vivo* expression measurements. We also use minireporters as small fusion tags to measure TXTL protein and peptide production yield. Finally, we generalize our concept by providing a luminescent minireporter based on split-nanoluciferase. The ∼80% gene sequence length reduction afforded by minireporters lowers synthesis costs and liberates space for testing larger devices while producing a reliable output. In the peptide production context, the small size of minireporters compared to full-length GFP is less likely to bias peptide solubility assays. We anticipate that minireporters will facilitate rapid and cost-efficient genetic device prototyping, protein production, and interaction assays.

## INTRODUCTION

Linear cell-free systems (CFS) are TXTL systems that enable rapid gene circuit characterization through the expression of linear DNAs, which eliminates the need for time-consuming cloning and large-scale plasmid preparation steps^1^. Linear DNAs are rapidly degraded by the RecBCD machinery in native *Escherichia coli* TXTL systems. Several methods have been developed to circumvent this limitation^1–4^, and to enable parts prototyping using linear DNAs. Coupling commercial DNA synthesis with linear TXTL prototyping offers a powerful and versatile workflow, supporting the rapid screening of DNA parts and their products. Linear dsDNA strands encoding the target genetic devices can be designed, synthesized, and directly tested upon reception, only requiring a PCR step when the synthesis scale is insufficient for directly using fragments out of the box (**Figure 1A**). As an example, we previously performed a full Design-Build-Test cycle in a week, the DNA part taking only a few hours^4^.

**Figure 1:**
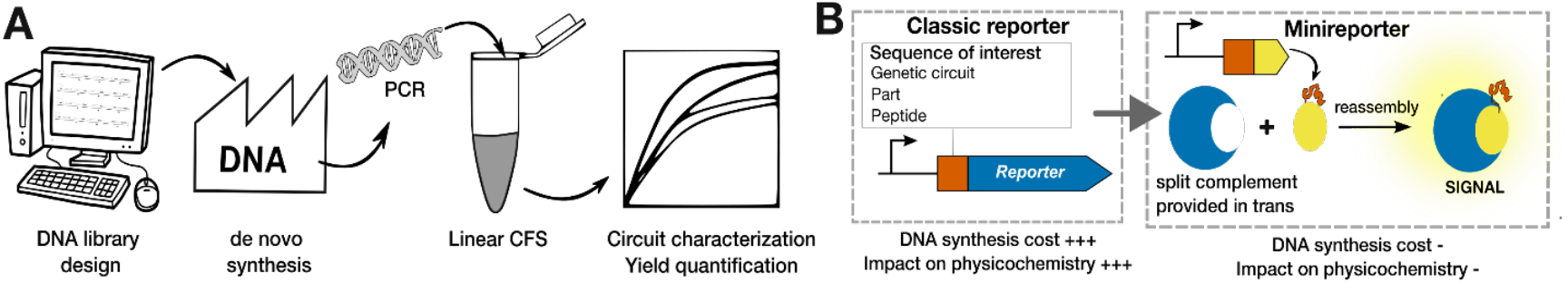
Workflow for fast prototyping using linear CFS and general principle of minireporters. **A**. Upon reception of short linear DNA fragments, a less than a day workflow enables characterization of the gene circuits and quantification of the produced peptide. **B**. Split reporters can be used as a smaller and cost efficient alternative for screening and prototyping libraries of genetic circuits and peptides.

As device characterization often requires the synthesis of a reporter protein (*e*.*g*. GFP or luciferase), screening various constructs in linear TXTL using total DNA synthesis requires repeated synthesis of the reporter gene, sometimes accounting for most of the synthetic DNA length and cost. In addition, rapid DNA synthesis workflows (i.e. 1-3 days) are generally available for limited DNA length, and the size of the test device is thus constrained by the reporter gene length. Possible alternatives include short fluorescent RNA aptamers^3,5^, but they have a low signal output and require the addition of costly fluorescent probes. Other systems based on DNA strand displacements that have been used in transcription-only CFS^6^ have a high signal-to-noise ratio, but their functionality in *E. coli* TXTL is unclear and they often require costly PAGE-purified DNA. All these systems enable fluorescence reporting of transcription only, and cannot be used to monitor protein or peptide production yields.

Here we sought to provide low-cost, robust, and protein-based alternative small reporters that would be functional in *E. coli* TXTL. Split-GFP systems, in which GFP is separated into two fragments that can reassemble when co-expressed, are ideal candidates. Initial split-FPs were used for Bimolecular Fluorescence complementation (BiFc)^7^, with two fragments of 154 AA and 83 AA (462 bp and 249 bp, respectively). Subsequent work using superfolder GFP (sfGFP)^8^ led to highly asymmetric split-GFPs in which a large barrel can reassemble with one or two small fragments from the short GFP C-terminus part. The first system developed^9^ is composed of a GFP1-10 large fragment (651 bp, 24.2 kDa) and a small GFP11 fragment (48 bp, 1.8 kDa). This system is routinely used to monitor protein solubility in living cells. Later on, a tripartite version was engineered^10^, in which the large GFP1-9 fragment (582 bp, ∼22 kDa) can reassemble with two small fragments derived from the two last C-terminal beta-sheets of GFP, GFP10, and GFP11, when those are in close proximity (*e*.*g*. when fused to interacting proteins). Interestingly, a bipartite system composed of GFP1-9, and a fusion between the two small fragments, termed GFP10-11 (135 bp, 5.2 kDa), also produces a functional GFP when reconstituted. We reasoned that these small GFP fragments could be exploited as short-length reporters (or *minireporters*). When synthesized, the minireporter would reassemble with its corresponding large fragment provided in *trans* (either from another linear DNA strand or expressed in *E. coli* before extract preparation) **(Figure 1B**). Minireporters lengths represent respectively ∼7% (GFP11) and ∼19% (GFP10-11) of the full-length GFP, a significant gain in space and synthesis costs for device characterization. We thus sought to optimize GFP11 and GFP10-11 minireporters in linear TXTL system.

## RESULTS

We first compared the fluorescence properties of different GFP reporters: sfGFP, and two GFP proteins corresponding to the fusion between the large and small fragments for GFP1-10 and GFP11 (termed FL11), and GFP1-9 and GFP10-11 (termed FL10-11) (**Figure 2A**). Genes encoding these proteins were placed under the control of the strong constitutive promoter P7^11^, self-cleaving ribozyme RiboJ^12^, and a strong bicistronic RBS (BCD2)^11^. We observed that sfGFP produced a much stronger fluorescence than FL11 (∼15% of sfGFP, 6 times lower) and FL10-11 (∼38% of sfGFP, 2.6 times lower). This result shows that the full-length protein FL10-11 arising from the reassembly of GFP1-9 and GFP10-11 has better fluorescent properties in TXTL than FL11. A more efficient complementation of the GFP1-9/10-11 couple compared to GFP1-10/11 was previously observed^13^. Regarding the smaller fluorescence intensity observed with both reconstituted proteins, these data could be explained by the fact that while the split variants were evolved from sfGFP, their reassembly occurs in a less sturdy barrel structure which results in reduced quantum yield and fluorescence brightness^14^.

**Figure 2:**
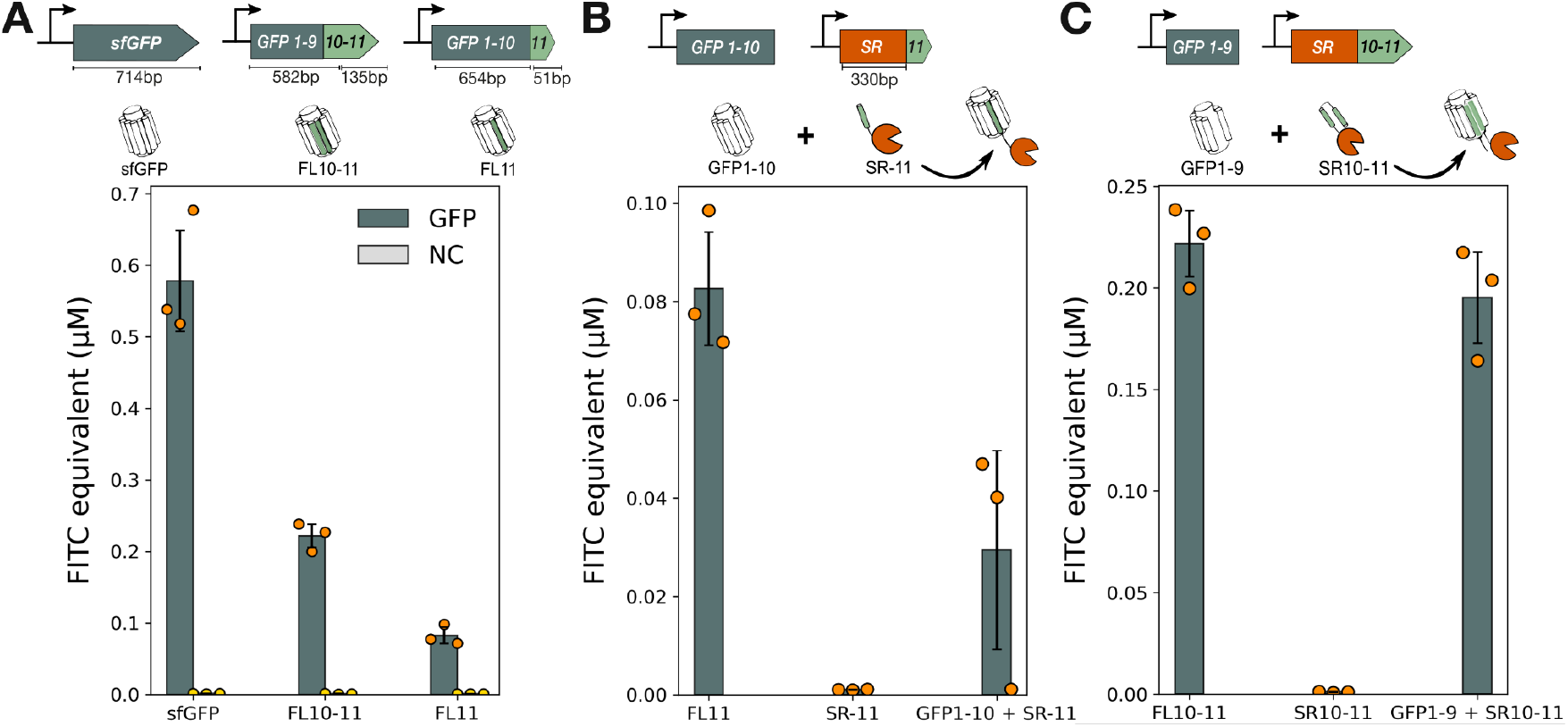
Functionality of split-GFP systems in linear CFS. **A**. expression of sfGFP, FL10-11 and FL11 in TXTL. NC: negative control, no DNA. **B**. Reassembly of SR-GFP11 with GFP1-10 in TXTL and comparison with FL11. **C**. Reassembly of SR-GFP10-11 with GFP1-9 in TXTL and comparison with FL10-11. All proteins were expressed from linear DNA at 5 nM concentration for each fragment and fluorescence values extracted at the 8-hour timepoint. Experiments were performed in triplicates on three different days. Error bars: +/- SD. See methods for more details.

Having established this approach, we co-expressed from two linear DNAs the large fragments (GFP1-10 and GFP1-9) along with their cognate small fragments fused to the sulfide reductase (SR11 and SR10-11), a canonical 110 AA protein used in split reporter assays^9^. We detected a fluorescent signal when the SR fusion was expressed with its corresponding GFP fragment, but none when expressed alone (**Figure 2B and 2C**). These data confirm that the reconstitution of split-GFP systems is possible in TXTL systems using linear DNA. Interestingly, fluorescence arising from the SR10-11 reconstitution was close to the level of its corresponding FL10-11 (∼95%). On the other hand, SR11 reconstitution, while producing a detectable signal, reached only 50% of FL11. Taken together, the SR experiments suggest that, at least in *E. coli* TXTL, fragment reconstitution is more efficient for the GFP10-11 system than for the GFP11 one.

To reduce reporter footprint, we then evaluated the most efficient split GFP to use as a stand-alone fluorescent reporter of gene expression. We expressed GFP11 and GFP10-11 using the same expression cassette with their corresponding split-GFP partner. As another minireporter version, we fused the GFP fragments to Tc5b (20 AA, 2 kDa, 60 bp), a synthetic, globular, and ultrastable Trp-cage small designer protein^15^. When coexpressed with GFP1-10, both GFP11 and Tc5b-GFP11 failed to produce a significant fluorescent signal **(Figure 3A**). This could be due to the poor reconstitution efficiency already observed with SR11. On the other hand, the GFP10-11 fragment produced a detectable signal when expressed with GFP1-9 **(Figure 3B**). This signal was about ∼30% of the fluorescent signal obtained with SR10-11. Interestingly, Tc5B-GFP10-11 produced a greater signal, close to SR10-11. This difference with GFP10-11 alone, together with GFP-11 data, suggests that smaller fragments might be more difficult to produce in TXTL systems or that they might be highly unstable. Also, synthesis of the larger GFP fragment from linear DNA might consume resources that could be used to produce larger amounts of the minireporter. In light of these results, we chose to use GFP10-11 as a minireporter for gene circuit prototyping.

**Figure 3:**
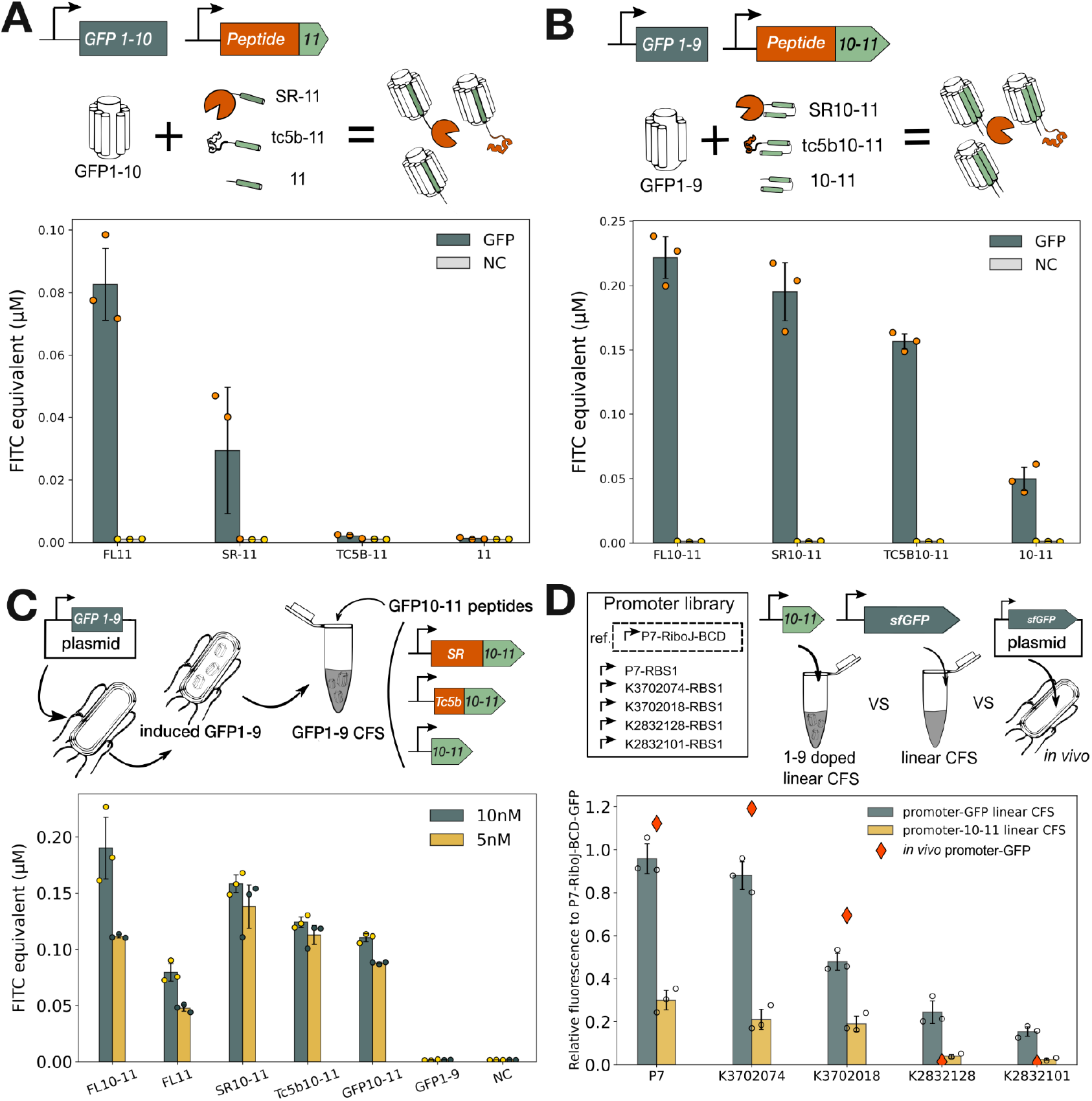
Optimization of minireporters based on split-GFP and application to promoter library screening. **A**. Comparison of the fluorescence intensity produced by FL11 with the coexpression of GFP1-10 with GFP11 fused to SR, Tc5B or alone. NC: negative control (extract without DNA). Fluorescence was extracted at 8 hours. Both linear DNA were added at a 5nM final concentration **B**. Same as in A, but comparing FL10-11 with the coexpression of GFP1-9 with GFP10-11 fused to SR, Tc5B or alone. **C**. Split-GFP reconstitution in extract doped with GFP1-9, and effect of linear DNA concentration on the measured fluorescence intensity, in comparison with full-length proteins. **D**. Assessment of promoter activity using the GFP10-11 minireporter, sfGFP, and comparison with *in vivo* activity. GFP1-9 doped extracts were supplemented with linear DNA fragment encoding different promoters driving expression of sfGFP (green bars) or GFP10-11 minireporter (gold bars). In parallel, the same constructs were cloned in a plasmid and their activity assessed in E. coli cells (red diamonds). All experiments performed in triplicates on different days. Error bars: +/-SD. See methods for more details.

### Cell-free Optimization of the GFP10-11 reporter in extracts containing the GFP1-9 fragment

To simplify the characterization process and maximize the resources available for genetic device activity, we prepared a cell-free extract from a strain containing a plasmid expressing the GFP1-9 fragment under an inducible T7 promoter (**Figure 3C**). As expected, this extract produced strong fluorescence when we expressed GFP10-11 alone or Tc5b-GFP10-11 from linear DNA. Interestingly, the fluorescence intensity measured from GFP10-11 was comparable to the one from the Tc5b fusion. One explanation is that the resources freed by the extract containing GFP1-9 enable greater expression of GFP10-11. Also, the presence of folded GFP1-9 in the extract might favor faster reassociation and stabilize GFP10-11. We tested two DNA concentrations, 5 nM and 10 nM, and found that the latter produced a greater signal for GFP10-11, but that the improvement was marginal for the Tc5b fusion. Of note, when using 5 nM DNA as in the previous experiment performed in non-doped extract, we observed a reduction of the reconstituted GFP protein signal. When we doubled the DNA concentration for reconstituted proteins (10 nM instead of 5 nM as in standard extracts), we could restore a signal close to the one observed in standard extracts. Because experiments from Fig 2 and 3 were done using different batches, it is hard to assess if this variation represents an intrinsic difference in extract protein production capacity or just a batch-to-batch variation. Despite those differences, we could use the doped extract to perform assays.

### Rapid assessment of promoter activity using split-GFP reporter and linear TXTL. Comparison with *in vivo*

Having obtained a reliable signal using the GFP10-11 fragment, we aimed to showcase the application of this minireporter system in linear TXTL by measuring the activity of a small set of promoters. We chose five promoters from the iGEM registry. We created three sets of constructs with a simple architecture - Promoter-RBS1-reporter. RBS1 (or UTR1) containing the T7 *g10* leader sequence is a strong consensus RBS from the classic pBest vector^16^. We built three sets of expression cassettes. In the first set, the five promoters drive the expression of GFP10-11. In the second set, the promoters drive the expression of sfGFP. The third set consists of the second set cloned in the pSB4K5 plasmid backbone for *in vivo* measurement (**Figure 3D**). A reference construct with known strong constitutive expression (P7-RiboJ-BCD-reporter), was also constructed in each set and used to normalize the fluorescence expression levels of the promoters set in each system. We measured fluorescence expression levels of the first and second sets in linear TXTL. We found that the rank order of promoters was comparable using these two reporters, validating GFP10-11 as a reliable minireporter protein (**Figure 3D**). Fluorescence intensity measured using GFP1-10 was smaller than with sfGFP, as already observed (Fig1A). Expression levels of the third set were measured *in vivo* by flow cytometry (**methods**). Relative promoter activities measured *in vivo* were comparable with rank order obtained in linear TXTL. These data suggest that linear TXTL using minireporters is a valid approach for prototyping genetic elements destined to be used in living cells.

### Split GFP-11 can be used to monitor protein production

While exploring the potentiality of GFP10-11 as a gene expression reporter, we investigated the use of GFP11 to monitor small peptide production yield, a current challenge in the TXTL field. Small peptides like antimicrobial peptides (AMP) are of interest but can be difficult to express or inherently toxic to the cellular host. While new methods for testing de novo AMP were shown to generate promising candidates against multidrug-resistant pathogens^17^, a fast and cost-efficient assay to determine peptide expression, folding, and concentration would be advantageous. TXTL was previously used to produce proteins and peptides of interest^18^. Quantifying protein synthesis and solubility using a fluorescent reporter would facilitate screening and optimization procedures. However full-length GFP tags increase the cost of *de novo* synthesis and likely interfere with self-assembly and solubility of the peptides, especially for small peptides consisting of a few dozen amino acids. Here we aimed to assess how the smallest split fragment GFP11 could be used as a reporter for peptide production and yield quantification *in vitro*. We reasoned that the fluorescence signal induced by the split assembly of a fused peptide-GFP11 could be a direct indicator of the peptide production yield in linear TXTL. Given that tc5b-GFP11 under a strong constitutive promoter did not produce a large signal, we turned toward a T7 expression system. We also assessed our fluorescence-based quantification method by comparing it to a standard His-Tag purification method. (**Figure 4A, methods**). The small fragment GFP11 did not produce a detectable signal when expressed as a standalone peptide in the presence of the GFP1-10 fragment. The GFP11 fragment was also not detected on a Tricine-based SDS-Page protein gel (**Figure 4C, methods**). We fused GFP11 to two mainstream peptides of different size: Sumo (103 AA, 12 kDa) and the TEV protease cleavage site (6 AA, ∼1 kDa). A 6-histidine tag was also added upstream to the peptide to allow his-tagged protein purification. We observed a strong fluorescence signal with 6His-SUMO-GFP11 and 6His-TEV-GFP11 site at 10 nM linear DNA concentration, while no signal was detected for GFP11 alone in the presence of GFP1-10 **(Figure 4B**).

**Figure 4:**
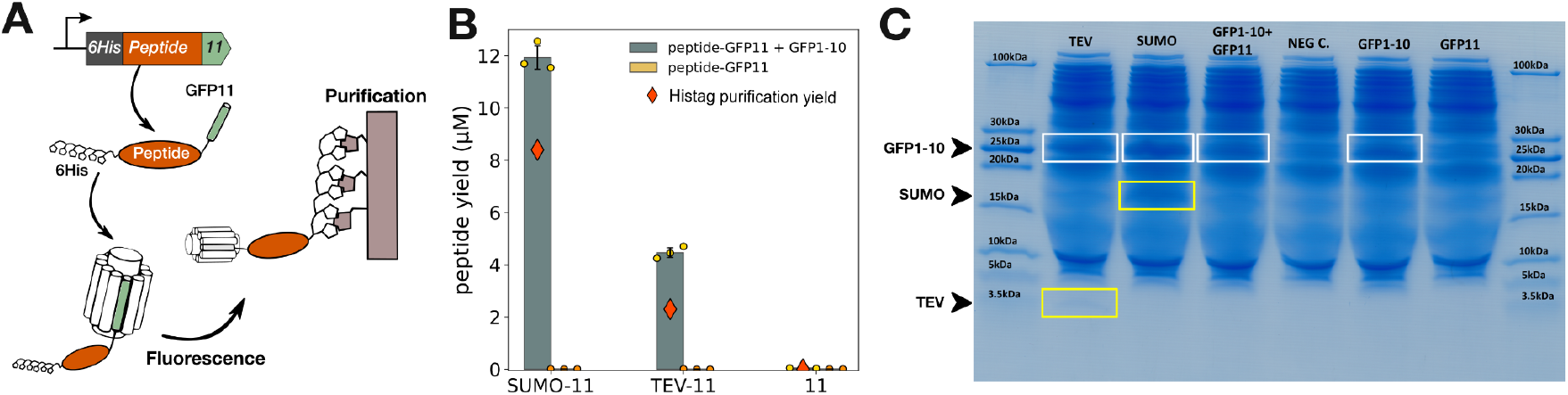
application of split-GFP minireporter to the monitoring short peptides production in linear CFS. **A**. Two different peptides, SUMO and TEV are fused to a 6-histidine tag and GFP11 (resp. N and C termini). Fluorescence was observed in linear CFS upon co-expression of the 6His-peptide-GFP11 with GFP1-10. His tag purification was performed sequentially to assess absolute peptide concentration. **B**. Gray bars show mean peptide concentration obtained by fluorescence intensity values from a microplate reader calibrated with purified GFP. Red diamonds show peptide concentration obtained after sequential His Tag purification and Bradford-based protein assay. **C**. Tricine-SDS-Page protein electrophoresis of 6His-peptide-GFP11 cell-free reactions. White frames highlight 6His-TEV-GFP11 and 6His-SUMO-GFP11. Black frames GFP1-10. Negative control corresponds to cell-free+water. No band was detected for GFP11 alone. The original un-labeled and uncropped protein gel is available in Supplementary material. See methods for more details.

By calibrating the plate reader with purified GFP, we estimated peptide titers to almost 12 µM for 6His-SUMO-GFP11 and slightly more than 4 µM for 6His-TEV-GFP11. His-Tag purification of the same samples was performed and yielded respectively around 8µM and 2 µM. These data indicate that the fluorescence-based assay could provide an accurate concentration of peptide produced in TXTL reaction. We further confirmed the presence of 6His-SUMO-GFP11 and 6His-TEV-GFP11 by protein electrophoresis directly performed on TXTL reactions (**Figure 4 C**). These data show that GFP11 can be used to monitor protein production efficiency in TXTL. In particular, short peptides could be produced and protein yield estimated via a simple fluorescent readout. This could be relevant for the production of antimicrobial peptides, in which the smaller (12 AA) GFP11 tag could have a lower impact on peptide activity than GFP10-11 (45 AA). Moreover, we show that peptides as small as 3 kDa can be expressed in a lysate-based TXTL solution from a linear DNA template on a tricine-based SDS-PAGE. This indicates that lysate-based TXTL is compatible with the expression of small peptides as long as they are soluble.

### Extending the minireporter family with luciferase derivatives

Finally, we sought to expand our approach to another type of minireporter based on a split nanoluc luciferase^19–21^. The split nanoluc system is composed of a small (smBit 33bp, 11 AA) and a large (LgBit 468bp, 156 AA) fragment (**Figure 5A**) that, unlike the split GFP system, do not spontaneously reassemble in solution. This can be of particular interest for protein-protein interaction studies. To promote the reassembly of the fragments in linear TXTL, we used a pair of antiparallel heterodimeric coiled-coils P3/AP4^22^. We fused P3 to the C-terminus of LgBit and AP4 to the N-terminal of smBit to respect the structural orientations of both fragments in the native nanoluc protein. To test this system in linear TXTL, LgBit, and smBit were introduced as linear DNA templates. As expected, individual fragments, and LgBit mixed with smBit did not produce any significant luminescence signal. On the contrary, LgBit-P3/AP4-smBit co-expression led to an increase in luminescence intensity by 7600-fold. (**Figure 5B**). Increasing the incubation time from 1 to 4 and 6 hours in linear TXTL before adding the luminescent substrate resulted in lower luminescent signals (**Figure S5**). This effect may be caused by the acidification that occurs over time in TXTL reactions^23,24^, and that was shown to affect nanoluc activity^25^.

**Figure 5:**
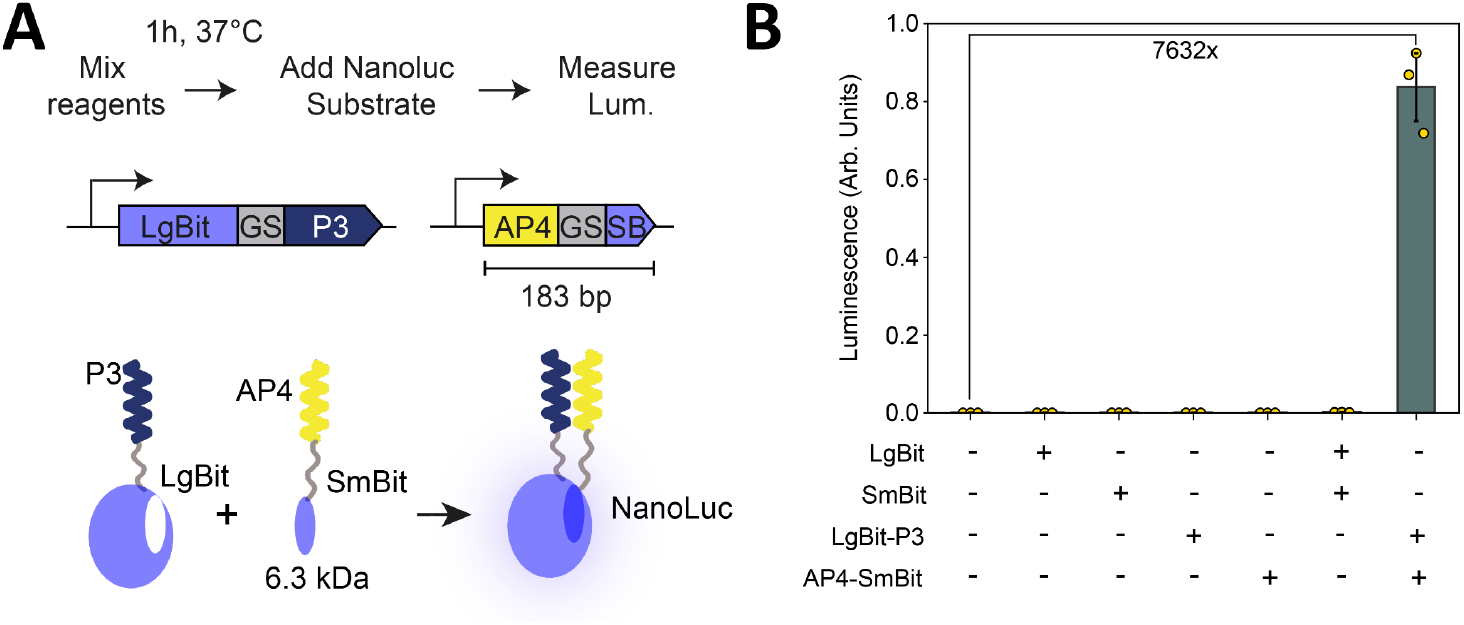
Functionality of minireporters based on split nanoluc in linear CFS. **A**. Schematics of the workflow and linear DNA parts used to reconstruct a functional Nanoluc from split LgBit and SmBit domains. **B**. Mean luminescent signal produced by adding the indicated constructs in linear TXTL, measured after 1h incubation. Error bars: +/-SD. See methods for more details.

These data demonstrate that minireporter can be declined to various systems, such as split enzymes, depending on the user’s need. In this case, the nanoluc reporter might provide, at the cost of adding a substrate and losing information about reaction kinetics, a better signal-to-noise ratio and limit of detection for experiments requiring higher sensitivity.

## DISCUSSION

Here we demonstrate that split-systems can be used reliably in linear TXTL systems to monitor genetic parts activity and peptide production. We tested a system based on GFP11 fragment and GFP10-11 fragment and found that the later provides a greater signal when used as a stand-alone reporter. While we did not recover a signal corresponding to the full-length protein when expressing both fragments from linear DNA, we were able to do so by doping the extract with GFP1-9 fragment, suggesting this issue arises from resource limitations in the context of coexpression of the two partners.

One limitation of our system is the lower signal intensity obtained when using split reporters compared to classical GFP assay. We observed that the fluorescent signal obtained from the full-length proteins corresponding to reassembled fragments is lower than the one obtained with sfGFP, using the same DNA concentration. While these variants could exhibit different transcription/translation rates or stability, previous study have shown that split-GFP variants have reduced quantum yield and fluorescence brightness due to a less sturdy barrel structure^14^. Future work might thus be directed at improving these properties in engineering new split variants. Nevertheless, the split GFP systems still provide a sufficient signal-to-noise ratio for most applications and luminescent minireporters can be used if needed.

GFP11 also proved a valuable tool to monitor and quantify peptide and protein production in TXTL. Some biases are possible in this system: the GFP11 tag which might improve peptide solubility and protein concentration. The split reporter could also impact the functionality of peptides that require self-assembly to be functional (such as some anti-microbial peptides). In that regard, split systems are not different from all other tagging systems that can have unintended effects, often as a function of the fusion protein sequence^26^. Yet, the power of simple, real-time measurement of peptide production yield in TXTL still represents a decisive advantage of the split system for rapidly screening peptide candidate libraries. A few best variants can then be chosen from a large set to undergo a more extensive characterization.

By lowering synthesis costs, allowing larger genetic devices to be prototyped, and protein production to be monitored in real-time, the minireporter systems presented here will expand the range of applications of linear *E. coli* TXTL.

## Authors contribution

AL, JB, SC, and VN designed the study. AL, JC, PM, PLV, ADV, performed the experiments. AZ, JB, and MCG supervised lab work at CBS. AL, JC, JB, and VN analyzed the results and wrote the paper.

## Associated content

Supporting Information is available.

- DNA sequences
- Original SDS-PAGE gel from Fig5
- Raw data and Jupyter Notebook used to analyze them are at: https://github.com/synthetic-biology-group-cbs-montpellier/Minireporters/tree/main

## Acknowledgments

We thank members of our groups and our institutes CBS for fruitful discussions and feedback. JB thanks INSERM and the Bettencourt-Schueller Foundation for continuous support. A.L. and I.K. gratefully acknowledges financial support for this publication by the Fulbright U.S. Student Program, which is sponsored by the U.S. Department of State, the Romanian-U.S. Fulbright Commission, and the Franco-American Fulbright Commission. A.L thanks Ecole Doctorale Frontières de l’Innovation en Recherche et Education – Programme Bettencourt and the Bettencourt Schueller Foundation for their generous support. The CBS acknowledges support from the French Infrastructure for Integrated Structural Biology (FRISBI) ANR-10-INSB-05-01. VN acknowledges support from the National Science Foundation (NSF CBET 2228971).

## Competing interests

The authors declare that no competing interests exist.

## METHODS

### Linear DNA preparation

DNA parts and constructs used in this study are listed in **Supplementary Table1**, and were ordered as g-blocks from IDT. The fragments were PCR amplified using primers listed in **Supplementary Table1** with Invitrogen Platinum SuperFi II, purified with Invitrogen PureLink Quick PCR purification kit, and eluted in nuclease-free water. PCR products were verified on a 1% agarose gel (1x TAE), aliquoted at 100 nM and stored at -20 °C.

### In vitro TXTL

Cell-free extracts were prepared according to a protocol previously published ^4,27^. Briefly, *E. coli* cells were grown in a rich medium, pellet, lysed with a cell press, clarified, pre-incubated, dialyzed, and stored at -80°C. A typical cell-free reaction contains *E. coli* lysate (33% v/v) supplemented with an energy mix, amino acids, and plasmids or linear DNA templates. Experiments were performed either using BL21 Rosetta2 extract supplemented with gamS^28^ to avoid linear DNA degradation (Fig 2, Fig 3A, 3B, 3C) or with BL21 Rosetta2 *ΔrecBCD* (Fig 3D, Fig4, and Fig 5).

### Fluorescence measurements

Quantitative measurements of gene expression were carried out using the reporter proteins deGFP and sfGFP. Fluorescence was monitored with a Cytation 3 Reader. 384-well square and covered by an adhesive plate seal (Thermo Scientific #AB0558) to minimize evaporation. Kinetic runs recorded fluorescence data at regular intervals for 8 h. End-point measurements were reported after 8 h of incubation at 37 °C. Raw data collected was converted to FITC Mean Equivalent Fluorescence (MEF) values for 20 μL reactions at 37 °C according to ^29^. FITC MEF and micromolar concentration values were obtained by calibration curves made using recombinant eGFP from ChromoTek GmbH (reference EGFP-250).

### In vivo promoter assembly and characterization

Promoter constructs were assembled in a pSB4K5 backbone by Gibson assembly ^30^ and cloned in DH5alphaZ1 ^31^. Assemblies were verified by Sanger sequencing (Eurofins, France). Plasmid sequences are available in supplementary data. For characterization, cells were grown 16 h at 37°C in Azure Hi-Def medium (Teknova, 3H5000) supplemented with 0.4% glycerol and kanamycin. For cytometry analysis, cells were diluted 200 times into Attune Focusing Fluid (Thermo Fisher Scientific, A-24904), and flow cytometry was performed on Attune NxT flow cytometer (Thermo Fisher) equipped with an autosampler and Attune NxT™ Version 2.7 Software. Experiments were performed in 96-well plates with settings; FSC: 200 V, SSC: 380 V, green intensity BL1: 460 V (488 nm laser and a 510/10 nm filter). All events were collected with a cutoff of 20,000 events. Every experiment included a positive control expressing GFP and a negative control harboring the plasmid but without a reporter gene. The cells were gated based on forward and side scatter graphs and events on single-cell gates were selected and analyzed, to remove debris from the analysis by Flow-Jo (Treestar, Inc) software. Geometric mean intensity values for three rounds of acquisitions performed on different days are reported.

### Protein electrophoresis

Protein electrophoresis was adapted after the method described by Schägger^32^. The need to use Tricine–SDS-PAGE compared to Glycine–SDS-PAGE arises as Tricine-based electrophoresis is able to resolve smaller size peptides (smaller than 30 kDa). The 16% Acrylamide AB-3 solution was used to cast a 0.7 mm 8x10 cm gel and run using the HoefHoefer™ Mighty Small™ II Mini Vertical Electrophoresis system. A 10-pin comb was used to create the wells. Prior to pouring the solutions for both the resolving gel and the stacking gel, the solutions were degassed for 10 minutes before the addition of APS and TEMED and degassed for a further 2 minutes after the addition of the two compounds. The loading samples were prepared by adding 3 uL of cell-free reaction, 12 uL of UltraPure water, and 5 uL of reducing loading buffer and incubated at 75 °C for 5 minutes. Each well was loaded with 7 uL of the prepared sample. The molecular weight markers used were the Thermo Scientific PageRuler™ Unstained Low Range Protein Ladder and Bio-Rad Precision Plus Protein™. The running conditions were constant voltage at 30 V for 30 minutes followed by constant voltage at 120 V for 1 hour and 10 minutes.

### His-tagged protein purification and Bradford assay

50 µL of cell-free reactions expressing 6his-peptides were pooled from 5x10 µL overnight reactions as starting protein extract material. His-tagged proteins were purified using 0.2 mL HisPur™ Ni-NTA Spin Columns from ThermoFisher according to the manufacturer’s protocol with the exception of carrying out 4 washing steps. Native conditions were used. The imidazole concentration of the Elution Buffer was chosen after eluting multiple samples with different concentrations of imidazole and measuring GFP fluorescence in the resulting elutions. The best imidazole concentration of the elution buffer used was 250 mM. After His-Tag purification the samples were subjected to protein quantification using the Bio-Rad Protein Assay Kit II and in a microplate reader (Synergy Neo2). The protocol recommended by the manufacturer was followed.

### Luciferase assay

10 nM of each linear fragment were mixed in a 22 ul TXTL reaction and incubated at 37 °C for 1, 4 or 6 hours. For luminescence measurements, 20 ul of linear TXTL reaction were mixed with 50 ul of Nano-Glo luminescent substrate (Promega N1110) into a white-bottom 96-well plate, and measured directly with a BioTek Synergy H1 Multimode Reader.

## Notes

https://github.com/synthetic-biology-group-cbs-montpellier/Minireporters/blob/main/Split%20minireporters.ipynb

### Competing Interest Statement

The authors have declared no competing interest.

